# Transactional sex among men who have sex with men participating in the CohMSM prospective cohort study in West Africa

**DOI:** 10.1101/630517

**Authors:** Cheick Haïballa Kounta, Luis Sagaon-Teyssier, Pierre-Julien Coulaud, Marion Mora, Gwenaelle Maradan, Michel Bourrelly, Abdoul Aziz keita, Stéphane-Alain Babo Yoro, Camille Anoma, Christian Coulibaly, Elias Ter Tiero Dah, Selom Agbomadji, Ephrem Mensah, Adeline Bernier, Clotilde Couderc, Bintou Dembélé Keita, Christian Laurent, Bruno Spire, the CohMSM Study Group

## Abstract

Men who have sex with men (MSM) are at much greater risk of HIV infection in Africa. Little is known about their involvement in transactional sex (TS). We aimed to characterize MSM reporting TS (MSM-TS) and to identify factors associated with their sexual practices using data from the prospective cohort study CohMSM conducted in Burkina Faso, Côte d’Ivoire, Mali and Togo. Our study focused on HIV-negative MSM, recruited between 06/2015 and 01/2018 by a team of trained peer educators. Scheduled study visits at 6, 12 and 18 months included medical examinations, HIV screening, risk-reduction counselling and face-to-face interviews to collect information on their sociodemographic characteristics, sexual behaviours, and HIV risk-reduction strategies. Three stigmatization sub-scores were constructed. The generalized estimating equation method was used for data analysis. Of the 630 HIV-negative participants recruited at baseline, 463, 410 and 244 had a follow-up visit at 6- and 12- and 18-months, respectively. Over a total of 1747 visits, 478 TS encounters were reported by 289 MSM-TS (45.9%). Of the latter, 91 participants reported systematic TS (31.5%), 55 (19.0 %) stopped reporting TS after baseline, and 53 (18.3%) reported TS after baseline. Ninety participants (31.1 %) reported occasional TS. After adjusting for country of study and follow-up visits, the following factors, reported for the previous 6 months, were associated with a greater likelihood of TS: younger age, an educational level <high-school diploma, satisfaction with current sex life, group sex with men, multiple male sexual partners, condomless anal sex, receptive or versatile anal sex with male sexual partners, giving benefits in exchange for sex with a man, alcohol consumption and drug use during sex, and experiencing stigmatization. The majority of MSM in this study who received benefits in exchange for sex had high-risk HIV infection exposure practices and were characterized by socioeconomic difficulties.

## Introduction

Men who have sex with men (MSM) are at a much greater risk of Human Immunodeficiency Virus (HIV) infection globally [1–3]. They constitute a highly heterogeneous group comprising bisexuals, transgender people and homosexuals from different socio-economic contexts and risk environments. These factors may influence their HIV exposure levels and their HIV transmission risk [4–7]. More specifically, age, educational level, economic disparities and income sources, housing stability, multiple sexual relationships and the exchange of benefits for sexual relations within these relationships (i.e., Transactional Sex, (TS)) have all been identified as factors related to different HIV exposure and transmission levels [8–10].

Studies have shown that MSM reporting TS (MSM-TS) are at a much greater risk of acquiring and transmitting HIV than MSM who do not report TS (MSM-NTS) because of high-risk sexual behaviours, including multiple sexual partners and frequent receptive anal intercourse. Various studies have also shown greater prevalence of HIV in MSM-TS than in MSM-NTS [11–17]. The increased vulnerability to HIV associated with TS is thought to be the result of multiple factors intervening at the individual (e.g., a large number of sexual partners, reduced negotiating power regarding the use of condoms, risky sexual behaviours because of alcohol and drugs, etc.) and structural (e.g., low socioeconomic status, limited access to healthcare and to stable housing, limited rights, and multi-layered social stigmatization, etc.) levels [18].

Literature defines TS as the exchange of gifts, money and/or other material benefits (food, shelter or other goods) for sexual activity, either regularly or occasionally. TS differs from commercial sex work in that sexual partners may also be boyfriends and girlfriends rather than simply clients and sex workers. In the African context, TS may be seen as part of a set of obligations for the boyfriend towards his girlfriend. In such relationships, this exchange is implicit; it is not formally negotiated and may not immediately follow a sexual act. More generally, shared emotional intimacy is present in many situations of TS [19,9,20]. The motivations to practice TS in men who receive benefits for sex (i.e., male sex workers) may be linked to needs related to poverty and wealth inequalities (‘survival sex’) or indeed needs related to a desire for trendiness, to popular culture, to the increased availability of commodities, and to the widespread use of global technologies (‘consumption sex’) [21,9,22].

Across sub-Saharan Africa, HIV disproportionately affects MSM compared with men in the general population [1,3,23,24]. While an overall decline in HIV prevalence has been noted in many geographic regions, HIV prevalence among MSM continues to rise (estimated in 2015 at 25.1 % in MSM-TS and 17.7 % in MSM-NTS [13,2]). Although various studies have highlighted that age, low educational level, condomless sex and alcohol use are all factors associated with TS and that they increase vulnerability to HIV among young girls in sub-Saharan Africa [20,25–28], little is known about these multilevel factors and the subsequent HIV risk among MSM-TS, especially concerning their social environment, motivation to practice TS and sexual risk behaviour. This is because they are usually included as subsets of larger studies focusing on MSM in general, and because MSM rarely report their TS practices and do not identify themselves as sex workers. Accordingly, this study was necessary to understand how to develop appropriate health education messages and tailored risk-reduction interventions for MSM-TS. Its aim was to characterize MSM who report TS in four West African countries (Burkina Faso, Côte d’Ivoire, Mali and Togo), and to identify factors associated with TS using follow-up data from a population-based prospective cohort study of MSM.

## Materials and Methods

### CohMSM Study procedures

In June 2015, a prospective cohort study of MSM was initiated at the premises of four local community-based organisations providing HIV services to MSM in four West African cities: Abidjan (Côte d’Ivoire), Bamako (Mali), Lomé (Togo) and Ouagadougou (Burkina Faso). Its main objectives were to assess both the feasibility and value of implementing a novel three-monthly preventive global care programme for MSM in West Africa, in order to help reduce the incidence of HIV in this key population, in their female partners and in the general population. The study did not compare a control group with an exposed group, nor was it based on a clinical trial. Potential participants were identified and recruited by a team of trained peer educators from the local organisations who approached individuals through a specific MSM network. Eligibility criteria included being at least 18 years old and reporting at least one anal sexual intercourse (insertive or receptive) with another man in the 3 months preceding study enrolment. Eligible individuals were offered a quarterly preventive global care package including: i) collection of information on health status, STI symptoms and sexual behaviours of individuals, ii) clinical examination, iii) diagnosis of STI and if necessary their treatment, iv) prevention tips tailored to MSM based on risk-reduction counselling, and v) provision of condoms and lubricants. In addition, vaccination against hepatitis B and annual tests for syphilis were proposed. HIV-negative MSM were also offered an HIV test at each quarterly visit. MSM found to be HIV-positive were offered immediate care for their infection, including ARV treatment. Before starting the interview, participants systematically received detailed information about the survey’s objectives and their right to interrupt the interview without justification. At enrolment and follow-up visits, participants completed face-to-face interviews with a research assistant who collected information on their sociodemographic and economic characteristics, HIV risk-reduction strategies, alcohol consumption, drug use and stigmatization. Participants had to provide written informed consent. The study team was very attentive to ensuring anonymity and confidentiality. Ethics approval was obtained from the National Ethics Committees of Mali (N°2015/32/CE/FMPOS), Burkina Faso (N°2015-3-037), Côte d’Ivoire (N°021/MSLS/CNER-dkn) and Togo (N°008/2016/MSPSCAB/SG/DPML/ CBRS). The study protocol was designed in accordance with the ethical charter for research in developing countries of French National Agency for Research on AIDS and Viral Hepatitis (ANRS) in France. The ClinicalTrials.gov Identifier is NCT02626286.

### Study population

Between 06/2015 and 01/2018, 778 participants were enrolled in CohMSM. As the study’s aim was to identify the risk exposure factors of HIV infection associated with TS, all HIV-positive participants (n=148) were excluded from the present analysis. Accordingly, this analysis focused on the remaining 630 HIV-negative MSM of whom 463, 410 and 244, respectively, had a follow-up visit at 6- and 12- and 18-months.

### Variables

#### Outcome

The study outcome was constructed on the basis of the following question: “During the last 6 months, have you been in a situation where you exchanged sex with a man in order to **receive** money, accommodation or any other benefit?”. This question was asked at baseline and all follow-up visits. Participants who responded “always” or “sometimes” at least once during the baseline and/or the follow-up, in contrast to those who responded “never”, were categorized as practising TS. The longitudinal nature of our outcome permitted us to identify participants who regularly practiced TS and those who intermittently practised it. Using this longitudinal approach ensured we did not lose any TS participants or related information about potential TS-associated variables which evolved over time.

#### Explanatory variables

a) Socio-demographic and economic characteristics: age was specified as a dichotomous variable dichotomised at the median (23.7 years). Categorical variables were constructed to indicate whether participants had at least a high-school level of education (=0 vs. < high-school=1 and not documented=2), were married or cohabitating (=0 vs. single, divorced or widowed=1 and not documented=2), and whether they had a stable housing status (=0 vs. unstable housing status=1 and not documented=2). Socio-economic characteristics included having an income generating activity (yes=0 vs. no=1), monthly income dichotomised at the median (50 000 Francs de la Communauté Financière en Afrique, approximately US$86.20 in 2019) and self-perceived financial situation (comfortable=0 vs. difficult=1 and not documented=2).

b) Sexual characteristics: a dichotomous variable indicated self-defined gender identity (man exclusively=0 vs. both a man and a woman=1); a self-defined sexual orientation identity variable indicated whether participants perceived themselves to be bisexual (=0 vs. not bisexual including homosexual, heterosexual=1); and a current sex life variable indicated the participant’s current level of sexual satisfaction (satisfactory=0 vs. very satisfactory=1). These variables are similar to those used in other studies [29–31]. A dichotomous avoidance variable was also constructed (no=0 vs. yes=1) to indicate whether participants practiced HIV risk-reduction strategies (e.g., avoiding sexual relations when drunk or when consuming other psychoactive products; using antiretroviral drugs to reduce the risk of HIV infection; avoiding anal penetration by seropositive partners or partners of unknown serostatus, etc.) (S1 appendix). Sexual behaviour was recorded using various variables: i) sexual position taken with male partners (exclusively insertive=0 vs. receptive or versatile=1 and not documented=2); ii) condom use with male partners during anal sex (yes=0 vs. no=1), iii) condom use with male partners during oral sex (yes=0 vs. no=1), iv) gel use with male partners during anal sex (yes=0 vs. no=1), v) disagreement about condom use with male partners (no=0 vs. yes=1), vi) number of male sexual partners (fewer than or equal to one=0 vs. more than one=1), vii) group sex with men (no=0 vs. yes=1), viii) use of psychoactive products during sex (alcohol no=0 vs. yes=1 and not documented=2, and drugs no=0 vs. yes=1 and not documented=2), ix) sudden sexual violence by male partners (no=0 vs. yes=1), and x) given benefits in exchange for sex with a man (no=0 vs. yes=1). The information provided by all these variables concerned the 6 months before the survey. Another variable, entitled “searching for male sexual partners on the internet” (no=0 vs. yes=1), concerned the previous 4 weeks. These HIV risk behaviour variables are similar to those used in other studies [32,4,15].

c) Stigmatization during the previous 6 months: the following three sub-scores, ranging from 0 to 10, were constructed, based on items from a previous study (S2 appendix) [33]: “experienced stigmatization during the previous 6 months” (based on 5 items, Cronbach’s alpha=0.83); “perceived stigmatization” (based on 11 items, Cronbach’s alpha=0.54); and “internalized stigmatization” (based on 8 items, Cronbach’s alpha=0.73).

Socio-demographic and economic characteristics were measured at baseline and were specified in the model as time-fixed variables. In contrast, sexual behaviour and stigmatization variables were measured at each time-point in the follow-up and consequently, were specified as time-varying.

### Statistical analysis

Descriptive analysis was conducted to compare baseline socio-demographic and economic characteristics, and sexual behaviours between MSM-TS and MSM-NTS. Categorical variables were compared between these groups using Pearson’s chi-squared test (χ2).

To identify factors associated with TS involvement, univariate and multivariate analyses were then performed using the generalized estimating equation (GEE) model which offers population-averaged estimates. A p-value < 0.2 was used to select variables for the multivariate model. The final multivariate model was estimated using a forward procedure. Fixed effects for each study country were specified in order to avoid bias arising from differences in sample sizes. All statistical analyses were performed using Stata version 13.0 (StataCorp, College Station, Texas, USA).

## Results

### Overall sample description

Over a total of 1747 visits, 478 TS encounters were reported by 289 MSM-TS (45.9% of 630 participants in the study). Of the latter, 91 MSM-TS reported regular TS (31.5%), 55 (19.0 %) stopped reporting TS after baseline, and 53 (18.3%) reported TS after baseline. Ninety MSM-TS (31.1 %) reported occasional TS.

**Table 1** shows the comparative analysis of baseline individual characteristics between MSM-TS and MSM-NTS. The former were significantly younger with a median age of 23.7 (p<0.001), and were significantly less likely to have a high-school diploma (27.7% *versus* 39.3%, p<0.001). A majority of MSM-TS (80.3% vs. 67.4 %, p=0.001) were significantly more likely to be unmarried (single, divorced or widowed). Almost a third of them had an income generating activity (28.4%), and 64.4% qualified their financial situation as difficult. Just over half (51.6%) had a monthly income lower than or equal to the median, corresponding to 50 000 FCFA. Despite having no work, MSM-TS were significantly more likely to have stable housing (68.2% increased probability, p=0.087). Moreover, 56.4 % of MSM-TS defined themselves as bisexual and were significantly more likely (47.9% increased probability, p=0.030) to define themselves as both a man and a woman. They were also significantly more likely (68.2% increased probability, p=0.008) to have receptive or versatile anal sex with male sexual partners (as opposed to exclusively insertive). Finally, they were significantly more likely to give benefits in exchange for sex with a man (15.2% increased probability, p=0.009).

**Table 1:** Comparative analysis of the baseline characteristics of the study sample (n=630)

| Sociodemographic and economic characteristics | MSM-TS<br>n=289 (45.9 %)<br>n (%) | MSM-NTS<br>n=341 (54.1 %)<br>n (%) | "p Value |
| --- | --- | --- | --- |
| Follow-up time-point |  |  |  |
| baseline | 199 (41.6) | 431 (34.0) | **<0.001** |
| 6 months | 96 (20.1) | 367 (28.9) |  |
| 12 months | 103 (21.6) | 307 (24.2) |  |
| 18 months | 80 (16.7) | 164 (12.9) |  |
| Study country (n=630) |  |  |  |
| Mali | 150 (51.9) | 99 (29.0) | **<0.001** |
| Cote d'Ivoire | 51 (17.7) | 84 (24.6) |  |
| Burkina | 41 (14.2) | 88 (25.8) |  |
| Togo | 47 (16.3) | 70 (20.5) |  |
| Age group relative to the median (23.7 years) (n=630) |  |  |  |
| >= the median | 114 (39.4) | 189 (55.4) | **<0.001** |
| < median | 175 (60.6) | 152 (44.6) |  |
| Educational level (n=630) |  |  |  |
| ≥ high-school diploma | 80 (27.7) | 134 (39.3) | **<0.001** |
| < high-school diploma | 182 (63.0) | 161 (47.2) |  |
| ND<sup>b</sup> | 27 (9.3) | 46 (13.5) |  |
| Marital status (n=630) |  |  |  |
| Married or living in a couple | 30 (10.4) | 65 (19.1) | **0.001** |
| Single, Divorced, Widowed | 232 (80.3) | 230 (67.4) |  |
| ND<sup>b</sup> | 27 (9.3) | 46 (13.5) |  |
| Had an income generating activity (n=630) |  |  |  |
| Yes | 82 (28.4) | 110 (32.3) | 0.291 |
| No | 207 (71.6) | 231 (67.7) |  |
| Monthly income relative to the median (50 000 Fcfa) (n=630) |  |  |  |
| <= median | 149 (51.6) | 165 (48.4) | 0.473 |
| > median | 106 (36.7) | 125 (36.7) |  |
| ND<sup>b</sup> | 34 (11.7) | 51 (14.9) |  |
| Financial perception (n=630) |  |  |  |
| Comfortable | 76 (26.3) | 103 (30.2) | 0.088 |
| Difficult | 186 (64.4) | 192 (56.3) |  |
| ND<sup>b</sup> | 27 (9.3) | 46 (13.5) |  |
| Stable housing (n=630) |  |  |  |
| Yes | 197 (68.2) | 205 (60.1) | 0.087 |
| No | 65 (22.5) | 90 (26.4) |  |
| ND<sup>b</sup> | 27 (9.3) | 46 (13.5) |  |
| Self-defined sexual identity (n=630) |  |  |  |
| Bisexual | 163 (56.4) | 184 (54.0) | 0.539 |
| Not bisexual | 126 (43.6) | 157 (46.0) |  |
| Self-defined gender identity (n=630) |  |  |  |
| Man exclusively | 150 (52.1) | 207 (60.7) | **0.030** |
| Both a man and woman | 138 (47.9) | 134 (39.3) |  |
| Sexual positioning with male partners in the previous 6 months (n=630) |  |  |  |
| Exclusively insertive | 85 (29.4) | 141 (41.4) | **0.008** |
| Receptive or versatile | 197 (68.2) | 192 (56.3) |  |
| ND<sup>b</sup> | 7 (2.4) | 8 (2.4) |  |
| Had given benefits in exchange for sex with a man (n=630) |  |  |  |
| No | 245 (84.8) | 312 (91.5) | **0.009** |
| Yes | 44 (15.2) | 29 (8.5) |  |

<sup>a</sup>p Calculated with Student's test
<sup>b</sup>ND = not documented. Includes missing data, “does not know” and “no response” terms.

### Factors associating MSM with Transactional Sex

Results from the multivariate analysis (**Table 2**) - after adjusting for the four study countries and follow-up time – indicated that the probability of practicing TS decreased by 68.9 % at 6 months of follow-up [adjusted odds ratio (aOR) and 95% confidence interval (95% CI):0.689(0.51-0.93)], although a tendency to increase by 39.5 % was estimated at 18 months of follow-up (aOR[95%CI]:1.395[0.98-1.98]). Younger participants were significantly more likely to practice TS (aOR[95% CI]:1.933[1.44-2.59]). In addition, having an educational level < high-school diploma (aOR[95%CI]:1.471[1.08-2.01]) was significantly associated with a greater likelihood of TS.

**Table 2:**
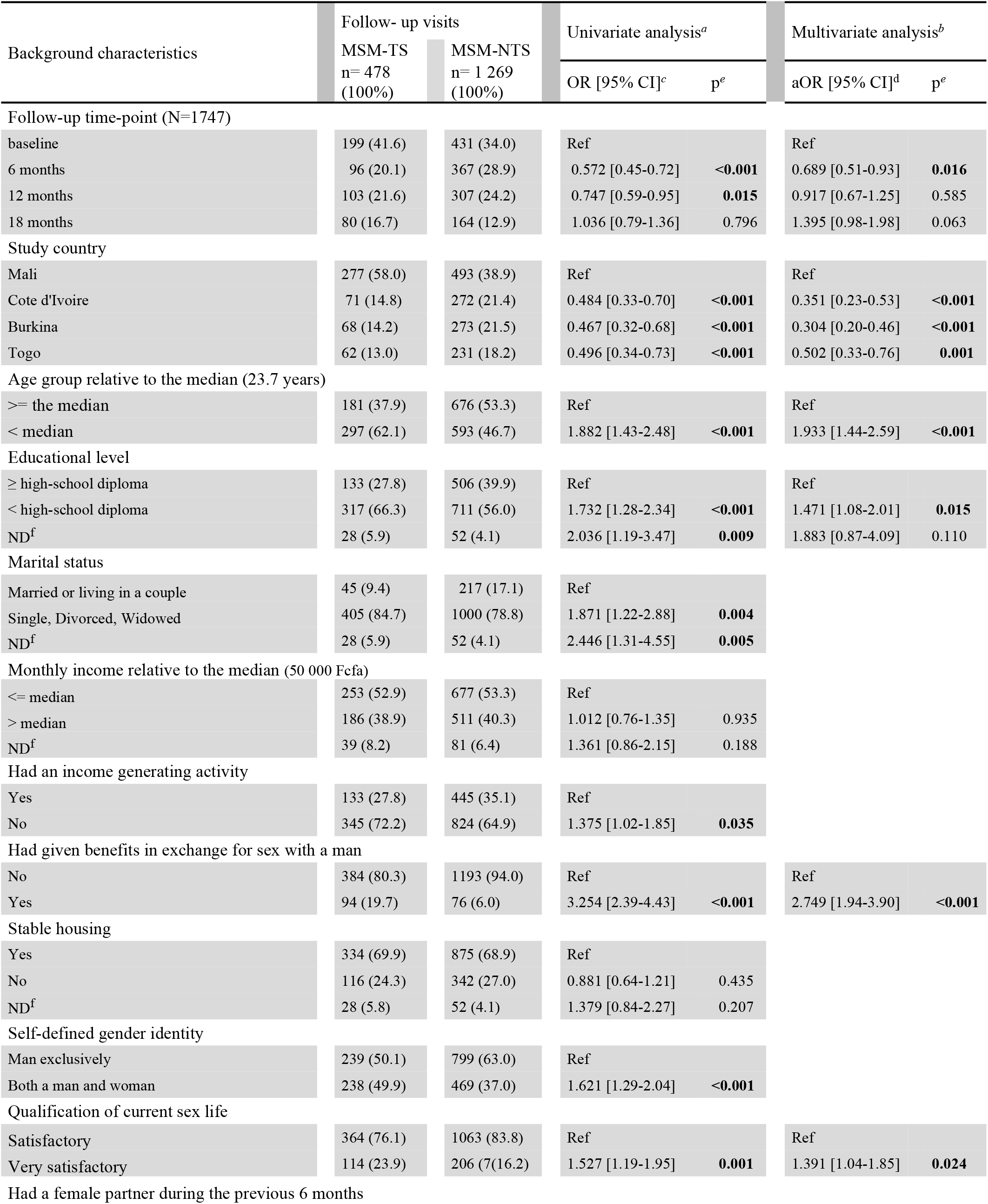
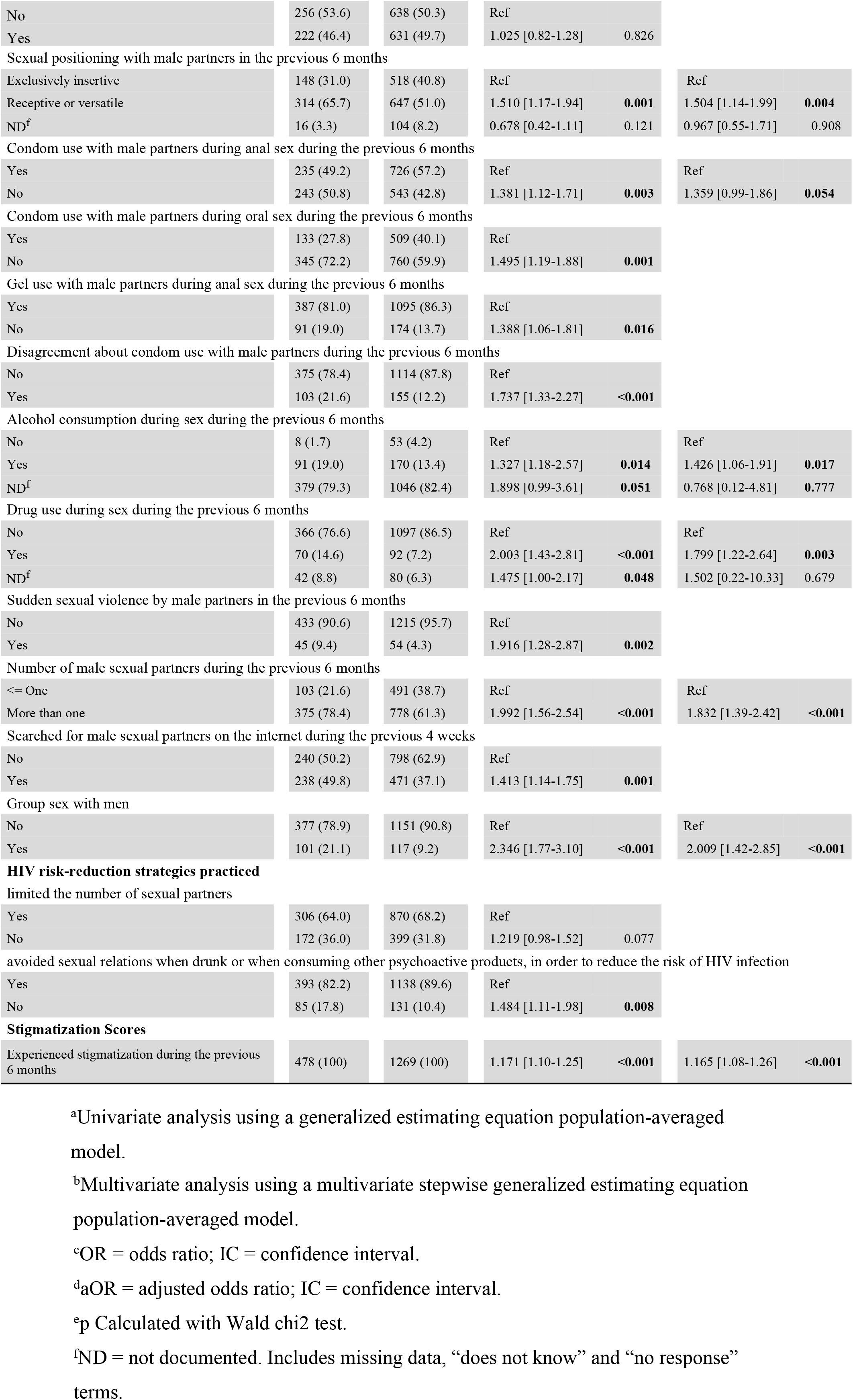
Factors associating Men who have sex with men (MSM) with transactional sex (TS) in West Africa: univariate and multivariate analyses with generalized estimating equation population-averaged model (n=630, 1747 follow-up visits)

With respect to sexual behaviours, TS was significantly more likely in those who had given benefits in exchange for sex with a man in the previous 6 months (aOR[95%CI]:2.749[1.94-3.90]), those who declared multiple male sexual partners in the previous 6 months (aOR[95%CI]:1.832[1.39-2.42]), and those who reported group sex with men (aOR[95%CI]:2.009[1.42-2.85]). Participants who self-reported practising receptive or versatile anal sex with male sexual partners in the previous 6 months were significantly also more likely to practice TS (aOR [95%CI]:1.504[1.14-1.99]). In addition, participants reporting any of the following factors were significantly more likely to engage in TS: a high level of satisfaction with their current sex life (aOR[95%CI]:1.391[1.04-1.85]), condomless anal sex in the previous 6 months (aOR[95%CI]:1.359[0.99-1.86]), alcohol consumption during sex (aOR[95%CI]:1.426[1.06-1.91]) and drug use during sex (aOR[95%CI]:1.799[1.22-2.64]) in the previous 6 months.

Finally, the more participants experienced stigmatization, the more likely they were to practice TS (this increase reaching 16.5%) (aOR [95%CI]:1.165[1.08-1.26]) during the previous 6 months.

## Discussion

Our results showed an overall rate of 45.9% for TS in MSM in the CohMSM cohort study. Those receiving benefits in exchange for sex had high-risk HIV infection exposure practices and were characterized by younger age, a low educational level, unmarried status and socioeconomic difficulties. Overall, our results for West Africa are in line with studies elsewhere (Tanzania, France), underlining a high percentage of MSM involvement in TS and an association between HIV risk factors and TS [4,15]. On the whole, we found a decreased probability of TS at 6 months of follow-up, but a tendency to increase at 18 months. As this was an interventional study where all MSM benefited from the same interventions, we are not able to explain these trends. In order to analyse the factors associated with TS, we were inspired by Baral et al.’s modified social ecological model (2013) [18]. Our results show that younger participants and those having a lower educational level were significantly more likely to practice TS. Although this finding contrasts with a study performed in several sub-Saharan African countries in 2005 which reported that educational level was not associated with engaging in TS among young men [34], it is in line with another study which analysed the associations between risky sexual behaviours, participation in TS, and HIV status among men and women aged 15-24 in Uganda [35]. These two factors - age and educational level – may increase HIV vulnerability in men, especially younger men with less sexual experience and men who have more difficulty accessing information on safe sexual behaviours in TS relationships [36]. Accordingly, strengthening education policy and improving school retention programmes for young people, especially MSM, would appear necessary. One possibility is to experiment with conditional cash transfer programmes. By improving young people’s economic situation, these programmes both help to retain students in school and incentivize safe sex. This leads to much less reliance on TS and therefore reduced risk [37].

Furthermore, our results highlight that MSM-TS were more likely to have condomless anal sex and to be very satisfied with their current sex life. In addition, MSM-TS were significantly more likely to have receptive sex or a versatile sexual position during anal intercourse. These practices (receptive sex and condomless anal sex) constitute the greatest risk of HIV transmission in MSM, particularly MSM-TS [38,39]. Our study did not collect data on the reasons for sexual satisfaction. However, another study showed that unprotected sexual intercourse is more satisfying, with MSM reporting that condom use increased sexual discomfort [40]. This may explain the high level of sexual satisfaction in our MSM-TS and the high level of reported condomless anal sex.

Our results also highlight that alcohol consumption and drug use during sex were associated with a greater probability of TS. There are multiple reasons why MSM-TS engage in substance use before and/or during sexual activity. Two of these are the burden of stigmatization of their sexual practices, and the search for strong sexual sensations [41]. These reasons may also be true for our study, but its design prevented us from being able to verify this. Furthermore, our results are in line with previous studies which highlighted that substance use among MSM-TS during sex increases the risk of HIV infection through increased risky sexual behaviours such as unprotected receptive anal sex [42–46]. This is also consistent with findings from studies in other sub-Saharan African contexts where substance use was associated with increased HIV sexual risk behaviours, which would in turn suggest an association between substance use and HIV prevalence and risk [47–49]. We believe that risk-reduction interventions focusing on substance use are necessary to mitigate the HIV epidemic in this population.

TS was significantly more likely to be reported by those declaring multiple male sexual partners during the previous 6 months, and those reporting group sex with men. This finding is consistent with other studies showing that the larger the sexual network of MSM, the greater the probability of engaging in TS with a member who already engages in it, as MSM tend to use their networks to find male sexual partners [50]. Consequently, there is a greater probability of being exposed to HIV-positive partners who do not practice HIV prevention measures [29]. Moreover, some studies have demonstrated multiple concurrent heterosexual partnerships and little or no condom use within transactional sexual partnerships in sub-Saharan Africa [51,25,9]. Hence, we recommend risk-reduction strategies that not only include components aimed at reducing multiple and concurrent sexual partners, but also include negotiation and communication skills aimed at encouraging systematic condom use among MSM. Furthermore, our results showed that monthly incomes of MSM-TS (86.20 US$) were very low compared with the GDP per capita (US$) of sub-Saharan Africa (3500 US$ for 2016). This confirms the low economic status of MSM in general and the wealth inequalities in their community [52]. In this context, young MSM may enter various sexual partnerships, often concurrently, because of multiple financial needs. It is therefore necessary to use a social and ecological model to understand MSM motivations for practicing TS, with a view to optimizing related social, behavioural and structural interventions.

Our results also showed that MSM-TS were significantly more likely to give benefits in exchange for sex with a man. This implies an overlap in the proportion of those who both received and gave benefits in exchange for sex with a man. These transactional sex dynamics contrast with prior research on MSM in Tanzania, Kenya and US [53,54,6], and seem to be based on a pattern wherein financial and material benefits play the role of a facilitator, with MSM receiving benefits for sex in situations where they would not by choice have sex with a particular partner, and giving benefits in situations where they have sex with a partner who does not find them attractive. Future studies, especially qualitative ones, should examine whether this group (i.e., receiving and giving benefits in exchange for sex with a man) has specific identifying characteristics or behavioral risks, in terms of sexual identity, partnership structure, and health. Knowledge of these characteristics and potentially associated risks could contribute to tailored health interventions for this group.

Finally, our study - which is the first to explore the different types of stigmatization (especially internalized) in MSM in West Africa – showed that MSM-TS were significantly more likely to have experienced stigmatization. To measure stigmatization, we decided to use the Homosexuality-Related Stigma Scale, developed and validated in Vietnam [33]. This scale takes into account all three types of MSM stigmatization (experienced, perceived and internalized). The simple questions used in the scale’s questionnaire allowed us to test for associations between TS and each type. Analyses yielded significant results, proving that stigma is a profound problem in this population. This confirms the value of performing more in-depth studies to validate this scale among MSM in West Africa. Social norms and the fear of being stigmatized may constitute barriers to finding regular sex partners, which in turn may push them to engage more in TS [55,7,56]. Although our results do not provide a reason as to why MSM-TS were more likely to be stigmatized than MSM not practising TS, it is possible that practicing TS with other men may lead MSM-TS to reveal their homosexual orientation more often when looking for a client, thereby increasing the risk of being stigmatized by the general public. Importantly, the high level of stigma experienced by MSM-TS may limit their use of healthcare services. This is in line with prior research which highlighted that access to and utilization of HIV prevention and care by the MSM population is influenced by certain social vulnerabilities such as stigmatization [29,57,58]. Our results suggest the need for stigmatization mitigation interventions to optimize MSM-TS linkage to HIV prevention and treatment services in West Africa.

The primary strengths of our study come from the fact that the CohMSM study was performed in four West African countries, was longitudinal in nature, and had four scheduled visits. Some study limitations should be taken into account when interpreting our results. First, we were not able to investigate our participants’ motivations for engaging in TS (i.e., out of financial necessity or for pleasure), or indeed whether they identified themselves as a sex worker or not. Second, given the declarative nature of the data and the fact that respondents participated in face-to-face interviews, social desirability bias is possible. Accordingly sexual risk behaviours may have been underreported. However, this bias was perhaps minimized by the fact that the research assistants involved all worked close to the ground, came from recognized non-governmental organizations, and were directly involved with the MSM population. Furthermore, participating MSM had a follow-up visit every 3 months. It is likely that a trustful relationship emerged over time with the research assistants, and consequently identifying whether the MSM engaged in TS became easier. Finally, we used stigmatization variables defined in the Homosexuality-Related Stigma Scale, developed and validated in Vietnam, but not used previously in West Africa.

Despite the study’s limitations, our results provide information useful for the optimization of prevention interventions for this at-risk high HIV prevalence MSM subgroup in West Africa. Biomedical, behavioural, and structural interventions such as the implementation of both early antiretroviral therapy and pre-exposure prophylaxis, as well as interventions to reduce stigmatization, are urgently needed to mitigate the effect of ongoing HIV transmission.

## Conclusion

Little is known about MSM-TS in West Africa. Our results show that the majority of MSM who received benefits in exchange for sex had high-risk HIV-infection exposure practices. Furthermore, this sub-group of MSM was characterized by younger age, a lower educational level, unmarried status and socioeconomic difficulties. Our results also show the importance of HIV prevention interventions among MSM-TS, and underline the need to develop more effective targeted prevention interventions at the community level (especially concerning the fight against stigma), as well as interventions addressing individual factors (pre-exposure prophylaxis, treatment as prevention and post-exposure prophylaxis). More in-depth multicentre research targeting MSM-TS is needed to better understand the multifaceted and multilevel factors associated with TS in West Africa, in order to take account behavioural heterogeneity in this population, and to clarify similarities and differences between countries.

## Supporting information

S1 appendix: Variables of HIV risk-reduction strategies. (S1 appendix.doc)

S2 appendix: Variables used to construct stigmatization scores. (S2 appendix.doc)

## Acknowledgements

**The CohMSM study group members are**: Clotilde COUDERC; IRD, INSERM, TransVIHMI, University of Montpellier, Bruno GRANOUILLAC; IRD, INSERM, TransVIHMI, University of Montpellier, Suzanne IZARD; IRD, INSERM, TransVIHMI, University of Montpellier, Christian LAURENT; IRD, INSERM, TransVIHMI, University of Montpellier, Laura MARCH; IRD, INSERM, TransVIHMI, University of Montpellier, Martine PEETERS; IRD, INSERM, TransVIHMI, University of Montpellier, Laetitia SERRANO; IRD, INSERM, TransVIHMI, University of Montpellier, Cheick Haïballa KOUNTA; Aix Marseille University, INSERM, IRD, SESSTIM, Cyril BERENGER; Aix Marseille University, INSERM, IRD, SESSTIM, Michel BOURRELLY; Aix Marseille University, INSERM, IRD, SESSTIM, Pierre-Julien COULAUD; Aix Marseille University, INSERM, IRD, SESSTIM, Gwenaëlle MARADAN; Aix Marseille University, INSERM, IRD, SESSTIM, Bakri M’MADI MRENDA; Aix Marseille University, INSERM, IRD, SESSTIM, Marion MORA; Aix Marseille University, INSERM, IRD, SESSTIM, Enzo PARISI; Aix Marseille University, INSERM, IRD, SESSTIM, Luis SAGAON-TEYSSIER; Aix Marseille University, INSERM, IRD, SESSTIM, Bruno SPIRE; Aix Marseille University, INSERM, IRD, SESSTIM, Adeline BERNIER; Coalition Internationale Sida, France, Paméla PALVADEAU; Coalition Internationale Sida, France, Daniela ROJAS CASTRO; Coalition Internationale Sida, France, Drissa CAMARA; ARCAD-SIDA, Mali, Oumar CISSE; ARCAD-SIDA, Mali, Alou COULIBALY; ARCAD-SIDA, Mali, Bintou DEMBELE KEITA; ARCAD-SIDA, Mali, Fodié DIALLO; ARCAD-SIDA, Mali, Mahamadou DIARRA; ARCAD-SIDA, Mali, Mady GADJIGO; ARCAD-SIDA, Mali, Abdoul Aziz KEITA; ARCAD-SIDA, Mali, Kader MAIGA; ARCAD-SIDA, Mali, Aly OUOLOGUEM; ARCAD-SIDA, Mali, Fodé TRAORE; ARCAD-SIDA, Mali, Niamkey Thomas AKA; Espace Confiance, Côte d’Ivoire, Camille ANOMA; Espace Confiance, Côte d’Ivoire, Stéphane-Alain BABO YORO; Espace Confiance, Côte d’Ivoire, Noufo Hamed COULIBALY; Espace Confiance, Côte d’Ivoire, Rachelle KOTCHI; Espace Confiance, Côte d’Ivoire, Patrick KOUABENAN; Espace Confiance, Côte d’Ivoire, Malan Jean-Baptiste KOUAME; Espace Confiance, Côte d’Ivoire, Kpassou Julien LOKROU; Espace Confiance, Côte d’Ivoire, Frédéric Dibi N’GUESSAN; Espace Confiance, Côte d’Ivoire, Xavier ANGLARET; PACCI, Côte d’Ivoire, Jean-Marie MASUMBUKO; PACCI, Côte d’Ivoire, Maxime OGA; PACCI, Côte d’Ivoire, Christian COULIBALY; Association African Solidarité of Burkina Faso, Ter Tiero Elias DAH; Association African Solidarité of Burkina Faso, Ousseni ILBOUDO; Association African Solidarité of Burkina Faso, Joseph OUEDRAOGO; Association African Solidarité of Burkina Faso, Mamadou OUEDRAOGO; Association African Solidarité of Burkina Faso, Elisabeth THIO; Association African Solidarité of Burkina Faso, Juste Rodrigue TOURE; Association African Solidarité of Burkina Faso, Abdoulazziz TRAORE; Association African Solidarité of Burkina Faso, Issa TRAORE; Association African Solidarité of Burkina Faso, Fiffou YOUGBARE; Centre National de Transfusion Sanguine, Burkina Faso, Nicolas MEDA; Centre de Recherche Internationale pour la Santé, Burkina Faso, Kouakou Kokouvi Selom AGBOMADJI; Espoir Vie Togo of Togo, Richard Mawuényégan Kouamivi AGBOYIBOR; Espoir Vie Togo of Togo, Messan ATTIOGBE; Espoir Vie Togo of Togo, Aléda Mawuli BADJASSIM; Espoir Vie Togo of Togo, Agbégnigan Lorette EKON; Espoir Vie Togo of Togo, Anouwarsadat KOKOUBA; Espoir Vie Togo of Togo, Ephrem MENSAH; Espoir Vie Togo of Togo, Diimiln Joseph Strauss TABLISSI; Espoir Vie Togo of Togo, Kossi Jeff YAKA; Espoir Vie Togo of Togo, Claver Anoumou Yaotsè DAGNRA; Laboratoire BIOLIM of the University of Lomé.

Lead author for CohMSM group: Christian LAURENT; IRD, INSERM, TransVIHMI, University of Montpellier, (**E-mail**)

